# Do inoculated microbial consortia perform better than single strains in living soil? A meta-analysis

**DOI:** 10.1101/2023.03.17.533112

**Authors:** Xipeng Liu, Siyu Mei, Joana Falcão Salles

## Abstract

Microbial consortium inoculation has been proposed as a natural-based strategy to safeguard multiple ecosystem services. Still, its empirical effects and comparisons to single-species inoculation have yet to be systematically quantified. In this global meta-analysis of 51 live-soil studies, we compared the impact (mean and variability) of single-species and consortium inoculations on biofertilization and bioremediation. Our results showed that both single-species and consortium inoculations increased plant growth by 29% and 48%, respectively, and pollution remediation by 48% and 80%, respectively, compared with non-inoculated treatments. We revealed the potential mechanisms contributing to the effectiveness of consortium inoculation, which are associated with the diversity of inoculants and the synergistic effect between frequently used inoculums (e.g., *Bacillus* and *Pseudomonas*). Despite a reduction in efficacy in field settings compared to greenhouse results, consortium inoculation had a more significant overall advantage under various conditions. We recommend increasing original soil organic matter, N, and P content and regulating soil pH to 6-7 to achieve a better inoculation effect. Overall, these findings support the use of microbial consortia for improved biofertilization and bioremediation in living soil and suggest perspectives for constructing and inoculating beneficial microbial consortia.

## 1. Introduction

Global concerns, such as increasing demands for food, disadvantages, and shortages of chemical fertilizers, and climate change, urged us to shift focus away from chemical inputs and towards sustainable techniques. Microbial inoculants are natural-based products that can potentially promote sustainability while safeguarding multiple ecosystem services (Eggermont et al., 2015; Alori and Babalola, 2018). Several globally accepted approaches focus on microbial inoculants showing potential for plant growth promotion, soil structure enhancement, and soil physiochemical properties optimization (Sharma et al., 2013; Mahanty et al., 2017; Santos et al., 2019). Additionally, taxa carrying out functions such as biosurfactant production, hydrocarbon degradation, and heavy metal accumulation are widely used in bioremediation (Zhuang et al., 2007; Fakhar et al., 2022; Wu et al., 2022).

Most commercially available products are based on single species, which successfully achieve the specific goal after validation, as evidenced by several recent meta-analyses (Thilakarathna and Raizada, 2017; Schütz et al., 2018; Tufail et al., 2021). However, individual species are limited in the functions they perform and the niches they occupy. This can potentially lead to intense competition between inoculant and native communities in the soil environment, resulting in significantly lower survival and functioning (Trabelsi and Mhamdi, 2013). Therefore, the quality and efficacy of microbial inoculants often show inconsistencies constrained by variable environments and application methods. This results in the limited practicability of microbial inoculants on a large scale (Alori and Babalola, 2018). Although these shortcomings can be partially compensated by combining other substances, such as chemical fertilizers, more sustainable methods are needed.

The idea of mixing multiple species with complementary traits, known as a consortium, was expanded in the last decade (Santos et al., 2019). The use of consortia is expected to obtain several advantages over single species. First, members in a consortium could compensate for traits that are lacking in other members to achieve a better effect. For instance, consortia consisting of isolates that produce indole-3-acetic acid and solubilize inorganic phosphates were more efficient at growth promotion than the strains applied individually (Kumar et al., 2016; Shilev et al., 2020). Other combinations could be plant-growth-promoting bacteria (PGPB) plus arbuscular mycorrhizal fungi (AMF) (Yadav et al., 2022) and drought-mitigating isolates plus nitrogen fixers (Prasanna Kumar et al., 2022), etc. Second, members in a consortium may facilitate the establishment and functioning of target strains due to the emergence of syntrophic cooperation between them (Sun et al., 2022). This interaction was also observed between PGPB and AMF, where plants could achieve a higher salt-stress resistance (Hashem et al., 2016) and better organic phosphorous mineralization (Jiang et al., 2021) compared to inoculated with either one of the microorganisms. Third, interactions between inoculants and indigenous species are more likely mediated through quorum sensing and antibiotic release within bacterial consortia (Santoyo et al., 2021). These aspects support the use of microbial consortia for a more stable and effective consequence (Aguilar-Paredes et al., 2020; Santoyo et al., 2021; Khan, 2022).

Despite these advantages, the application of microbial consortia is still disputed, with similar disadvantages as single species (Kaminsky et al., 2019; Jack et al., 2021). For instance, microbial inoculation represents a deliberate biotic disturbance (also known as microbial invasion) for the native soil microbiome, which can induce complex interactions between inoculated and native species, leading to unpredictable and unreliable outcomes (Liu et al., 2022). Besides, both the characteristics of consortium (e.g., composition, diversity, and density) and application strategies (e.g., inoculating in the soil or on the seed and plant types) regulate the inoculation effect. However, most microbial inoculants’ development is based on laboratory screening and combination. Whether and how to make microbial inoculants survive and function stably in the natural environment is still challenging (Kaminsky et al., 2019). Therefore, there is an urgent need to systematically and quantitatively investigate the efficacy of microbial consortium in living soil and explore the factors that may determine the inoculation effect to guide the subsequent development and application of microbial inoculants.

Here, we performed a systematic meta-analysis on data from peer-reviewed publications that compared the effects between single-species and consortium inoculations on biofertilization and bioremediation. Only experiments conducted in living soils were included (see “Materials and methods” for study selection details) to obtain quantitative results close to natural conditions. We identified the overall impact of using single-species and consortium inoculants and explored multiple factors that may influence the outcome of consortium inoculations. We hypothesize that (i) the inoculation effect of consortium in living soil would be better than single species based on the potential advantages mentioned above, (ii) environmental conditions and practical factors may influence the beneficial effect of inoculants while using consortia might be more reliable compared to single species, and (iii) the synergistic effect of members within the consortium contributes to the better inoculation effect.

## 2. Materials and methods

### 2.1. Literature search

We focused on peer-reviewed studies that measured bacterial inoculants’ effects within the setting of living soil that compared single inoculants and consortia. Searches were performed in Web of Science (Core collection) following guidelines from PRISMA (Preferred Reporting Items for Systematic Reviews and Meta-Analyses, Supplementary Fig. S1). Search terms were used in Topic: (soil AND (microbial OR bacterial) AND (consortia OR consortium OR communit*) AND (inocul*)). We included all articles published between January 2012 and December 2021, which resulted in 2149 studies.

Study titles and abstracts were scanned manually during the first selection, excluding studies that lacked information on the potential application of microbial inoculants in the soil. The second selection was performed by screening full texts to determine whether the studies met the following criteria: (1) a comparison performed between the bacterial consortium and single inoculant involved in this consortium, (2) an experiment conducted in living soils that harbored native microbial communities, (3) attributes (mean, standard error/deviation, and statistics) that could be extracted directly from the text, tables, and Figures. Finally, a total of 51 studies met the selection criteria and were used in the analysis. Locations of these studies and their trend across publication years were shown in Fig. 1 and Supplementary Fig. S2, respectively.

**Fig. 1.**
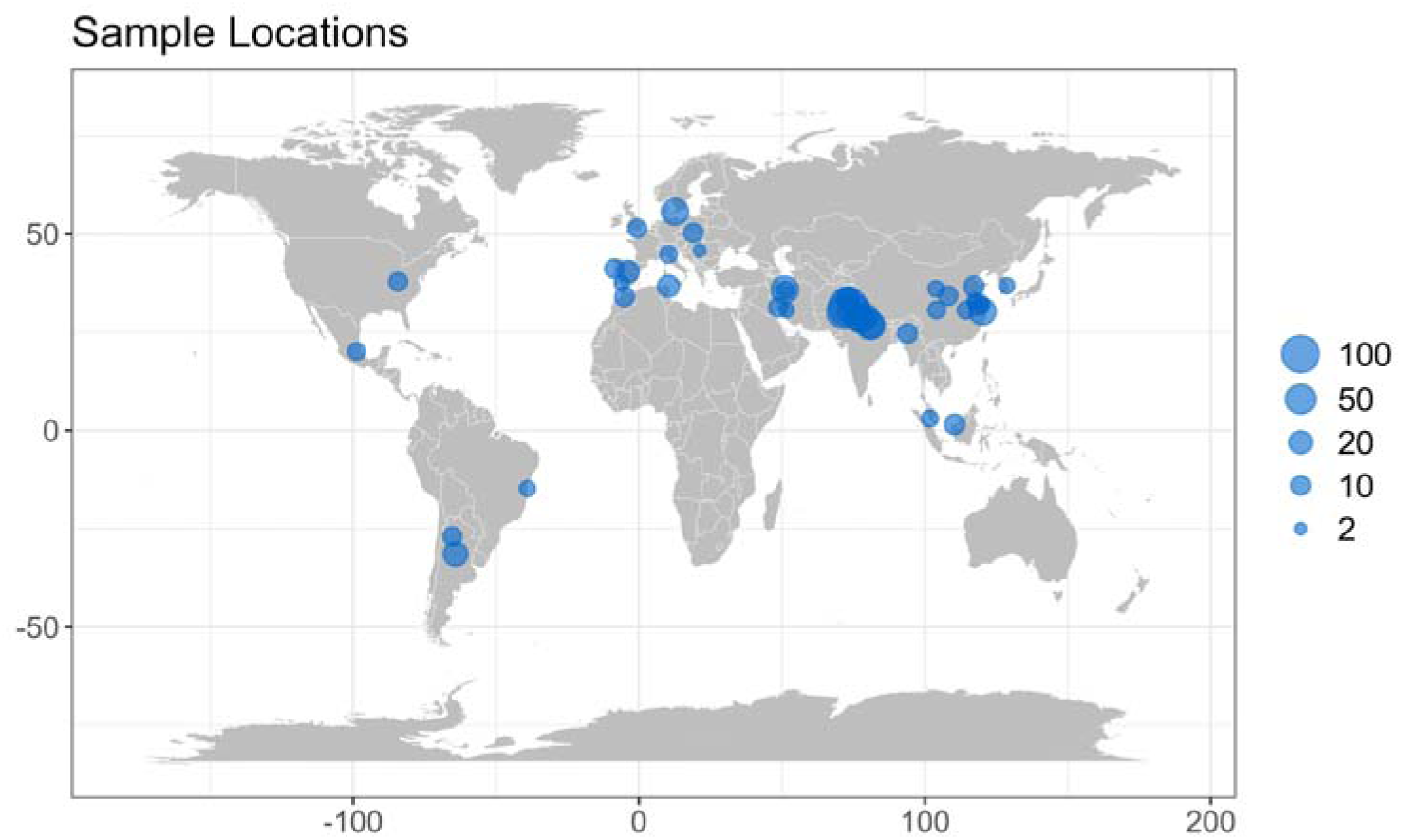
Global distribution of studies. 796 effect sizes (lnRR) from 51 studies were included in the meta-analyses. The size of the dots represents the number of effect sizes.

### 2.2. Screening and extraction

For each study, we extracted the mean, the number of replications, and the standard deviation of each indicator representing two main beneficial functions: biofertilization and bioremediation. To quantify the biofertilization effect, we extracted indicators related to plant growth, including dry biomass, yield, and seed germination. Other surrogate indicators, such as fresh biomass and plant height, were extracted when none of the above indicators were reported. For bioremediation, we extracted the heavy metal content in plant tissues or the degradation/removal rate of heavy metals or organic pollutants from soil. When both metal content in plant tissues and removal rate from the soil were provided, the former was recorded to prevent redundancy. When the information was presented only in graphical form, the data were digitized from Figures using Getdata Graph Digitizer (Version 2.26). The recorded standard error (SE) for each observed value was converted to standard deviation (SD) by the equation: 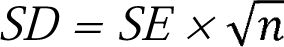, where *n* is the number of replications. For the studies only reporting the levels of statistical significance with letters, we use the TINV function in Excel to calculate the *t* value and convert it to *SE* by the equation: *SE = MD / t*, where the *MD* is the difference in means.

We obtained inoculant diversity (i.e., number of species), geographical location, experiment type, duration, soil properties, inoculation method, and plant/pollutant types. The experiment type indicated whether the experiment was conducted in soil microcosm (without plants), pot/greenhouse (with plants), and field (with plants) conditions. The inoculation time described the duration of the experiment, from inoculant application to sample collection. As one study did not provide the experiment duration, we used the mean of all studies to replace it. Inoculation methods included soil and seed inoculation, which consists of the application of microbial inoculants either directly by pouring into the soil or soaking the seeds/seedlings. The plant type was grouped into cereal and non-cereal plants, and the pollutant type was divided into heavy metal and organic pollutants. The properties of soil used in each experiment, including pH, texture (proportion of clay), and contents of organic matter (OM), nitrogen (N), and phosphorus (P) were extracted.

### 2.3. Effect sizes

Effect size calculations were completed using the *escalc* function in the R package ‘metafor’ (Version 3.8.1) (Viechtbauer, 2022). For the mean differences, we used a natural log-transformed response ratio (lnRR) as the effect size to compare inoculation effects in non-inoculated and inoculated systems. The formula is lnRR = ln (X_t_/X_c_), where X_t_ and X_c_ are the mean values in the treatment applied with single inoculant or consortium and without inocula, respectively. To quantify the effect of inoculations on the relative variability of different beneficial functions, we calculated the natural logarithm of the ratio between the coefficients of variation (lnCVR) using lnCVR = ln(CV_t_/CV_c_), where CV is SE/X, X is the mean value, SE is the standard error, t is the treatment applied with single inoculant or consortium, and c is the treatment without inoculum, respectively (Nakagawa et al., 2015). We obtained 796 effect sizes for both lnRR and lnCVR. The data set focusing on biofertilization consisted of 586 effect sizes from 45 studies, and on bioremediation consisted of 210 effect sizes from 15 studies.

### 2.4. Meta-analyses

The meta-analytic (intercept) model was fitted to assess the overall effect of inoculation in the absence of moderators. We then used inoculant type (i.e., single inoculant and consortium) as the main moderator to compare the difference between single inoculant and consortium in effect sizes. A meta-regression model was performed to investigate whether the inoculant species of consortium and inoculation duration affect the inoculant functioning. Pairwise comparisons of subgroups were conducted to assess the moderating effects of experiment types, inoculation methods, and plant/pollutant types. Statistical significance compared to the reference was assumed when 95% confidence intervals (95% CIs) did not span zero. The differences between subgroups were estimated by Tukey’s HSD comparisons using the *ghlt* function in “multcomp” package (Hothorn et al., 2008). The heterogeneity in each model was assessed using I^2^ (Higgins et al., 2003) and provided in Supplementary Tables. The index R^2^, describing the ratio of explained variance to the total variance, was calculated for each model using the *r2_ml* from the “orchard” package (Nakagawa et al., 2021) and provided in Supplementary Tables.

All models were run with the chosen random effect structure using the R package “metafor” (Version 3.8.1) for biofertilization and bioremediation data separately. The “orchard” package (Nakagawa et al., 2021) was used for visualization. The following sources of non-independence were identified and considered: species effects (e.g., inoculants within experimental cases and studies), case effects (e.g., experimental cases within studies), study effects (effect sizes calculated from the same survey), pairwise comparisons for effect size calculations (effect size ID; variability in the true effects within studies). The final random effect structure was set as study ID, case ID, species ID, and effect size ID after comparing AICc values among models.

For testing publication bias in our intercept models, we used the funnel plot based on the meta-regression residuals (Doleman et al., 2020) and the conventional Egger’s regression test (Sterne et al., 2001). The former was performed by fitting standard error as a unique moderator (Nakagawa et al., 2022). The results of publication bias are provided in Supplementary Fig. S3 and Table S1. We also tested the time-lag bias using meta-regression analysis to assess the relationship between publication year and effect size (Supplementary Fig. S4 and Table S2). All analyses were conducted using R (version 4.2.1).

## 3. Results

Our dataset includes globally distributed empirical studies conducted in living soil. These studies were conducted in Asia, Europe, North Africa, North America, and South America (Fig. 1, 51 studies and 796 effect sizes). Most studies focused on inoculation effects on biofertilization (N = 586, 73.6%), with the remaining on bioremediation (N = 210, 26.4%). By using the non-inoculated treatment as the reference, the counts of effect size were 358 (45.0%) and 228 (28.6%) for biofertilization and 149 (18.7%) and 61 (7.7%) for bioremediation under the single-species inoculant and consortium treatments, respectively.

### 3.1. Mean effect sizes of bacterial inoculations for biofertilization and bioremediation

Bacterial inoculants had an overall positive effect size than non-inoculated treatments (lnRR = 0.31, 36%, 95% CI = 0.24-0.37, *p* < 0.001; Fig. 2A and Table S3) for biofertilization function, which suggests the plant growth could be improved on average by 36% by using inoculants. The beneficial effect was also observed for bioremediation with an effect size of 0.46 (58%, 95% CI = 0.28-0.63, p < 0.001) compared with the control, non-inoculated treatment (Fig. 2B).

**Fig. 2.**
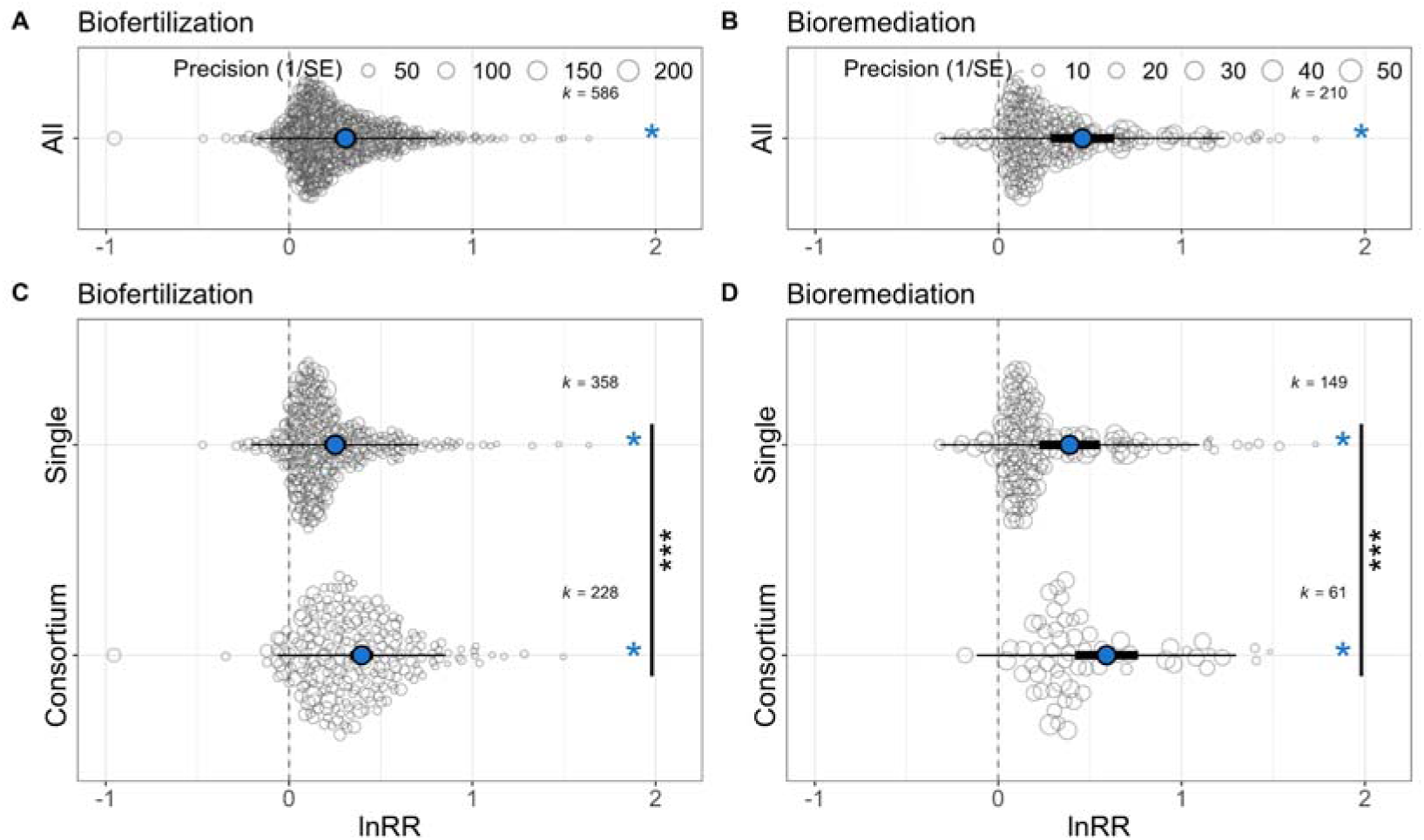
The beneficial effect of bacterial inoculants on biofertilization and bioremediation. The overall effects (A, B) and comparisons between single-species and consortium inoculants (C, D) were estimated. Blue points in the central represent estimated means, thick bars represent 95% confidence intervals, and thin bars represent 95% prediction intervals. Each background point is an effect size (lnRR), and its size is scaled by the precision of that estimate (1/SE). Blue asterisks denote the rejection of the hypothesis that the estimated mean equals 0 (*p* < 0.001), indicating a significant effect compared to non-inoculant treatments. Black asterisks suggest a significant difference in estimated means between the two groups (***, Tukey’s HSD, *p* < 0.001).

Subgroup analysis showed that both single-species inoculant and consortium treatments significantly affected biofertilization and bioremediation (Fig. 2C and 2D, Table S4). Applying single species showed an increased effect on plant growth promotion on average by 29% (lnRR = 0.25, 95% CI = 0.19-0.31, Fig. 2C and Table S4). Consortium applications had a significantly larger estimated effect size for biofertilization (lnRR = 0.40, 48%, 95% CI = 0.33-0.46) than single-species treatments (z value = 8.8, *p* < 0.001). For bioremediation, the increasing inoculation effect of the consortium was 80% (lnRR = 0.59, 95% CI = 0.42-0.76) and significantly higher than that of single-species treatments (lnRR = 0.39, 48%, 95% CI = 0.22-0.55, Fig. 2D and Table S4).

### 3.2. Variability of inoculation effects on biofertilization and bioremediation

Variation in inoculation impacts (lnCVR) was significantly decreased (less than 0) for biofertilization by 15% after bacterial inoculations, with the control, non-inoculated treatment (lnCVR = -0.16, *p* < 0.01, Fig. S5 and Table S5). Both applying single-species (lnCVR = -0.17, *p* < 0.01) and consortium (lnCVR = -0.15, *p* < 0.01) decreased the variation of inoculation impacts on biofertilization, but no significant difference was observed between them (Fig. S5). For bioremediation, no significant difference in variability was observed compared to non-inoculation treatments (Fig. S5, Table S5 and S6).

### 3.3. Moderators influencing the inoculation effect

For biofertilization, the application of bacterial consortium had a higher positive impact on plant growth, whether tested in pot/greenhouse (by 51.7%) or field (by 37.7%) conditions, compared to treatments applied with single-species inoculum (by 30.4% and 21.1% for pot and field settings, respectively) (Fig. 3A and Table S7). The estimated lnRR under pot/greenhouse and field conditions inoculants showed no statistical difference for single species (*p* > 0.05). Still, they were significantly different for the consortium, and the lnRR was lower under field conditions (lnRR = 0.42 and 0.32 for pot and field settings, respectively, *p* < 0.05) (Fig. 3A and Table S7). In contrast, no significant difference was observed for bioremediation between experiment types (i.e., microcosm and pot/greenhouse) (*p* > 0.05, Fig. 3B and Table S8). We found the lnCVR was significantly less than 0 only in pot/greenhouse conditions despite using single-species and consortium for biofertilization (*p* < 0.05, Fig. 3C), indicating that the beneficial effect of inoculation on decreasing the variability of plant growth mainly occurs in controlled environments. No significant impact was observed on lnCVR for bioremediation (*p* > 0.05, Fig. 3D).

**Fig. 3.**
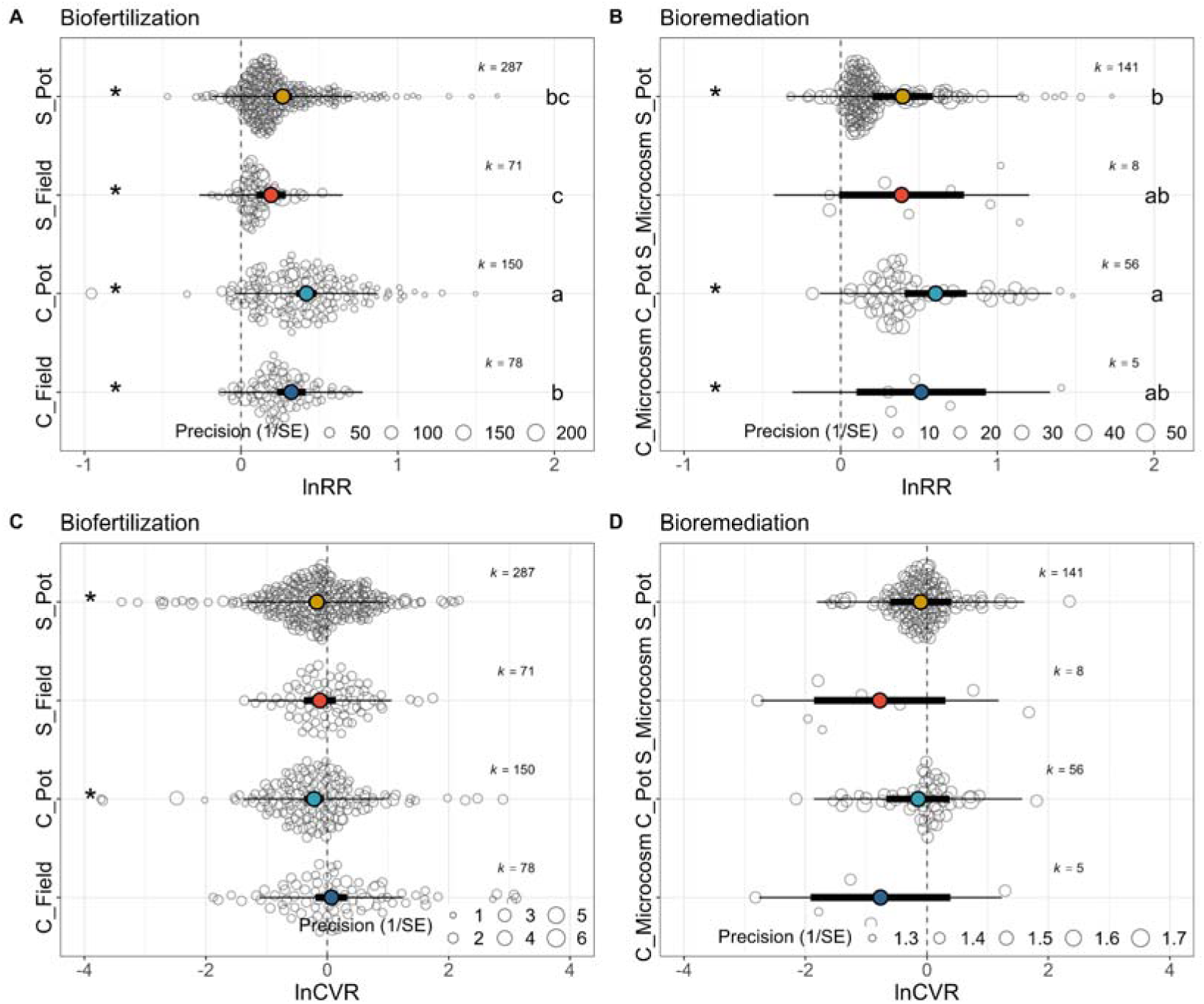
Impact of experiment types on effect sizes (lnRR) and variability (lnCVR) of biofertilization (A and C) and bioremediation (B and D). The experiment type indicates whether the experiment was conducted in soil microcosm (without plants), pot/greenhouse (with plants), or field (with plants) conditions. Letters S and C for subgroups represent single-species and consortium inoculations, respectively. Central points represent estimated means, thick bars represent 95% confidence intervals, and thin bars represent 95% prediction intervals. Asterisks represent the rejection of the hypothesis that the estimated mean equals 0 (*p* < 0.001), indicating a significant effect compared to non-inoculant treatments. Different letters on the right suggest a significant difference between subgroups (*p* < 0.05, Tukey’s HSD).

The inoculation method (i.e., soil inoculation and seed/seedling inoculation) had no significant influence on the estimated lnRR applying either single-species or consortium inoculum for biofertilization and bioremediation (*p* > 0.05, Fig. 4A and 4B, Table S9 and S10). However, a larger mean effect size was observed in consortium application treatment than single species treatment for biofertilization and bioremediation, regardless of inoculation methods (Fig. 4A and 4B).

**Fig. 4.**
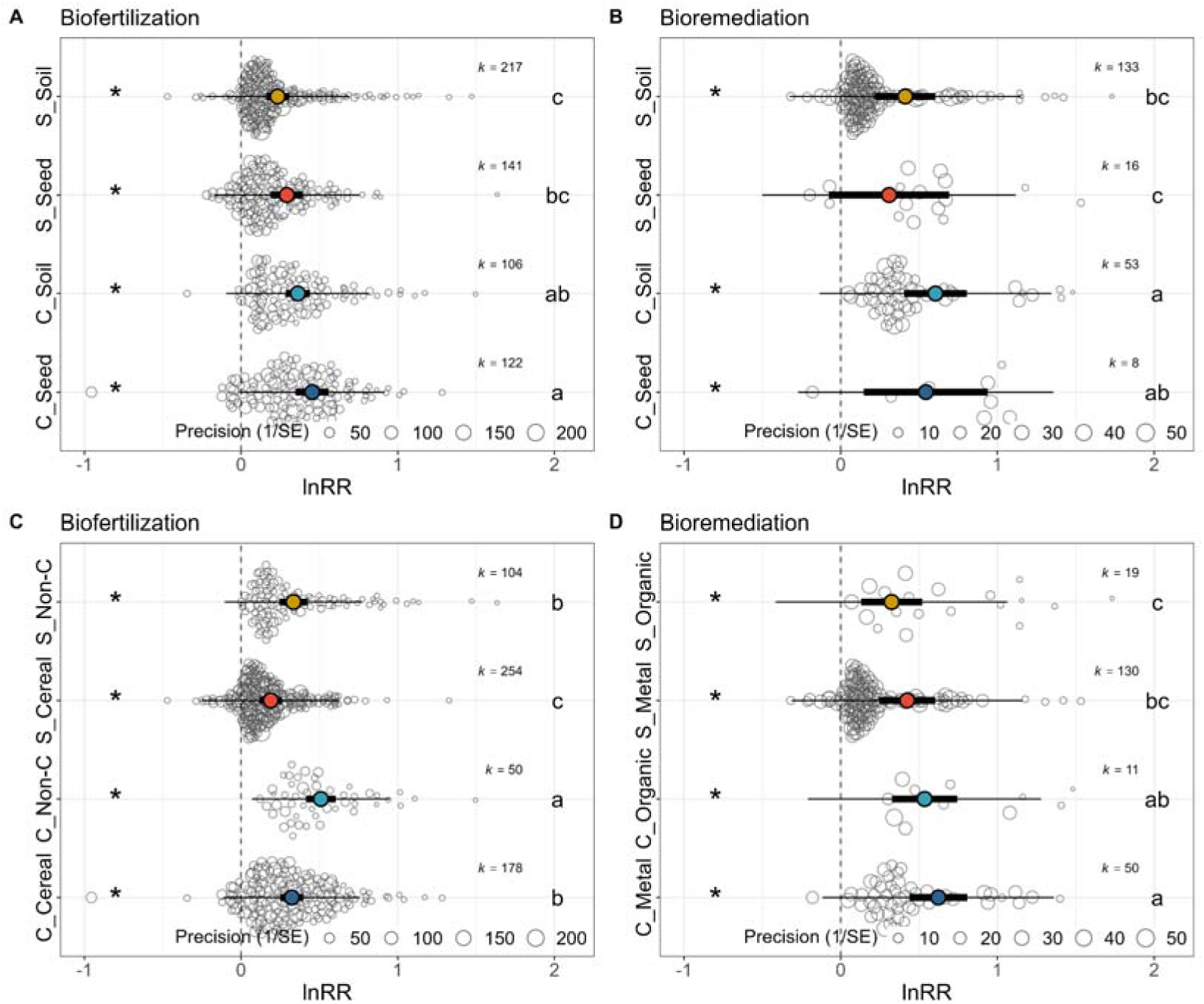
Impact of inoculation methods (A, B) and target types (C, D) on the inoculation effect of biofertilization and bioremediation functions. The plant type was grouped into cereal and non-cereal (Non-C) plants, and the pollutant type was divided into heavy metal and organic pollutants. Letters S and C for subgroups represent single-species and consortium inoculations, respectively. Asterisks denote the rejection of the hypothesis that the estimated mean equals 0 (*p* < 0.001), indicating a significant effect compared to non-inoculant treatments. Different letters on the right suggest significant differences between subgroups (*p* < 0.05, Tukey’s HSD).

Plant type was evaluated as an essential moderator of the inoculum’s function for biofertilization (Fig. 4C). Specifically, the lower estimated lnRR of cereal plants (lnRR = 0.19 and 0.32) was found compared to non-cereal plants (lnRR = 0.33 and 0.51) for single-species inoculations and consortium inoculations, respectively (*p* < 0.05, Fig. 4C and Table S11). For bioremediation, the effect sizes showed no difference between pollutant types (i.e., metal and organic pollutants) for both single-species inoculations and consortium inoculations (*p* > 0.05, Fig. 4D and Table S12). Furthermore, the advantage of applying consortium was observed for both biofertilization and bioremediation, regardless of plant or pollutant types (Fig. 4 and Table S12).

The impact of inoculation time (i.e., experimental duration) on effect sizes was further investigated. Our result showed that experimental duration had no significant correlation with the effect size for biofertilization, although the estimate was higher after consortium applications (*p* > 0.05, Fig. 5 and Table S13). For bioremediation, a weak impact on inoculation effects after the one-off inoculation was observed for both single-species (estimate = 0.0005, 95% CI = -0.0003-0.0014, t value = 2.11, *p* = 0.027) and consortium (estimate = 0.0017, 95% CI = 0.0003-0.0032, t value = 2.23, *p* = 0.021) treatments (Fig. 5 and Table S13).

**Fig. 5.**
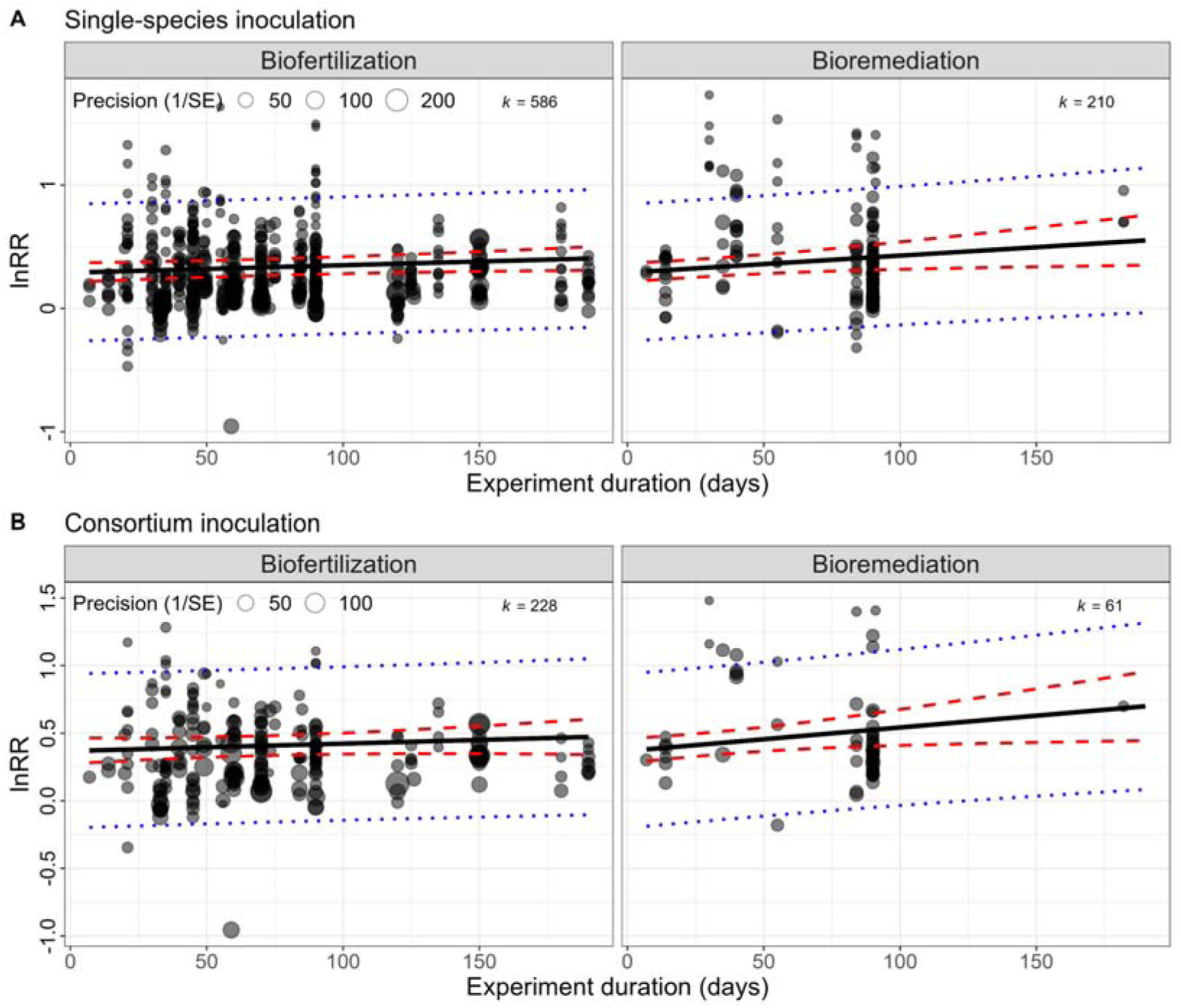
Meta-regression analyses assessing the impact of experiment duration on inoculation effects. The experiment duration had a weak impact on inoculation effects after the one-off inoculation (estimates = 0.0003, 0.0013, 0.0005, and 0.0017, *p*-value = 0.33, 0.22, 0.03, 0.02 for single-species and consortium inoculations for biofertilization and bioremediation, respectively). The red and blue dashed lines represent 95% confidence and 95% credibility intervals, respectively.

### 3.4. The influence of original soil properties on effect sizes

The properties of the soil used for inoculation experiments affected inoculation effectiveness (Fig. 6). The results showed that the relationship between soil pH and lnRR was significantly fitted with a polynomial curve (*p* = 0.0031) in the consortium treatment for biofertilization, but not in the single-strains treatment (*p* = 0.35). For bioremediation, original soil pH was negatively correlated with lnRR for both single-strain and consortium treatments (*p* = 0.00055 and 0.05, respectively). The content of organic matter in soil was significantly and positively correlated with lnRR in two treatments for biofertilization (*p* < 0.01). However, these relationships were not significant for bioremediation (*p* > 0.05).

**Fig. 6.**
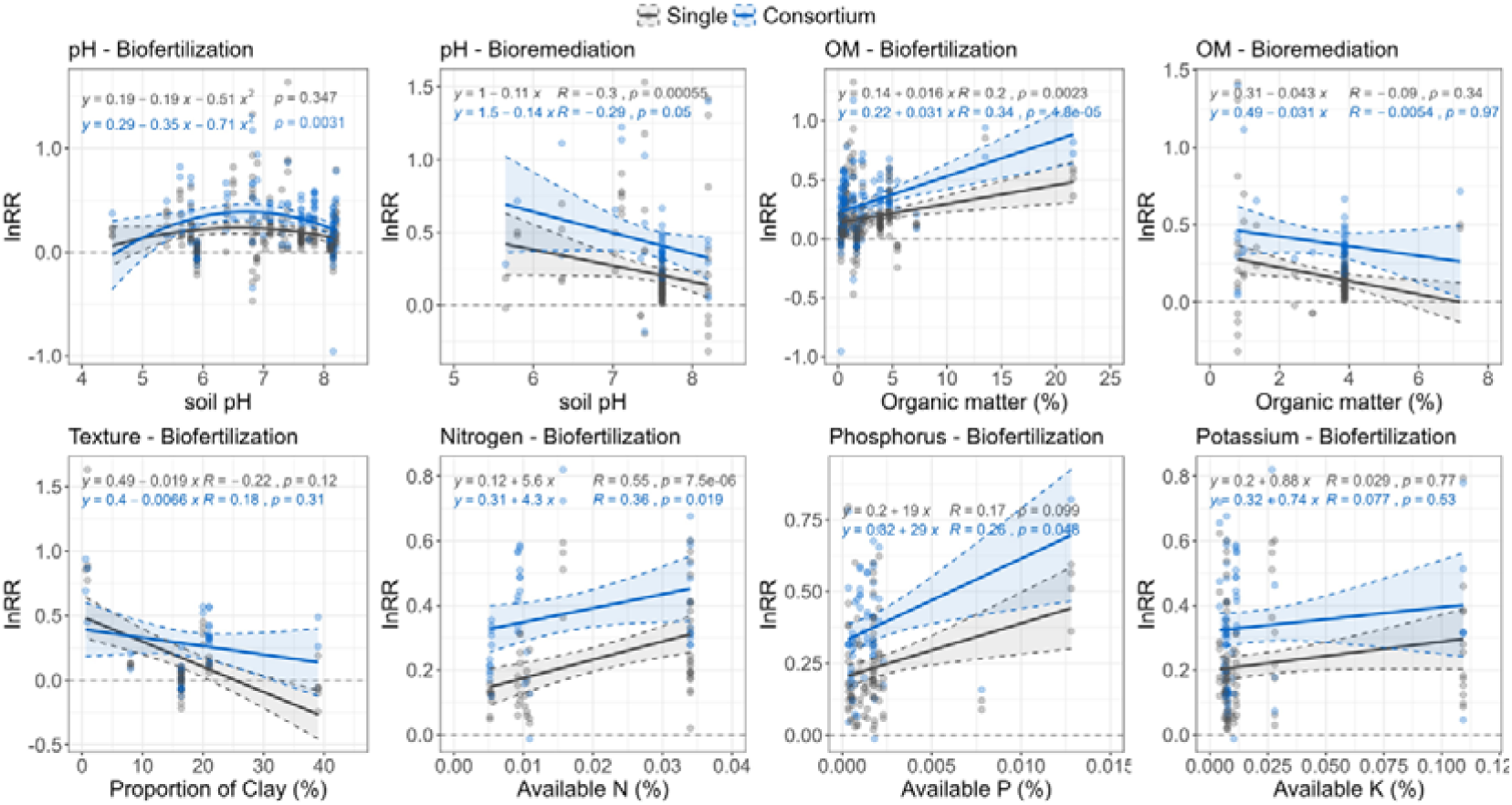
Impacts of original soil properties on inoculation effect. Relationships between effect sizes and soil pH, texture, and the content of organic matter (OM), available nutrients (N, P, and K), were estimated by polynomial curve fitting or linear regression (Spearman correlation).

The impacts of soil texture and available nutrients, such as nitrogen (N), phosphorus (P), and potassium (K), on effect sizes, were assessed only for biofertilization, since limited data was obtained from bioremediation cases (Fig. 6). No significant correlation between clay proportion and available K content in soil and lnRR was observed in neither single-strain nor consortium treatments for biofertilization (*p* > 0.05). Available N content in original soil was significantly correlated with lnRR in single-strain (R = 0.55, *p* < 0.0001) and consortium (R = 0.36, *p* = 0.019) for biofertilization. For available P content in soil, such significant and positive correlation was only observed in consortium treatment (R = 0.26, *p* < 0.048).

### 3.5. The influence of inoculant diversity on effect sizes

Meta-regression analyses identified the impacts of alpha diversity (number of species) of the inoculum and experiment duration (from inoculation to sample collection) on effect sizes. The number of species of the inoculum was a critical modulator influencing inoculation effects (F_2,793_ = 53.40, *p* < 0.0001, Fig. 7A and Table S14). For the biofertilization impact, the model’s estimated regression weight for inoculum species is 0.080 (CI = 0.062-0.099, 8%, t value = 8.73, *p* < 0.0001). This means that for every additional species, the inoculation effect of a study was expected to rise by 8%. For the bioremediation effect, the significant regression coefficient (estimate = 0.122, 13%, CI = 0.090-0.153, t value = 7.68, *p* < 0.0001) was observed across inoculum diversity.

**Fig. 7.**
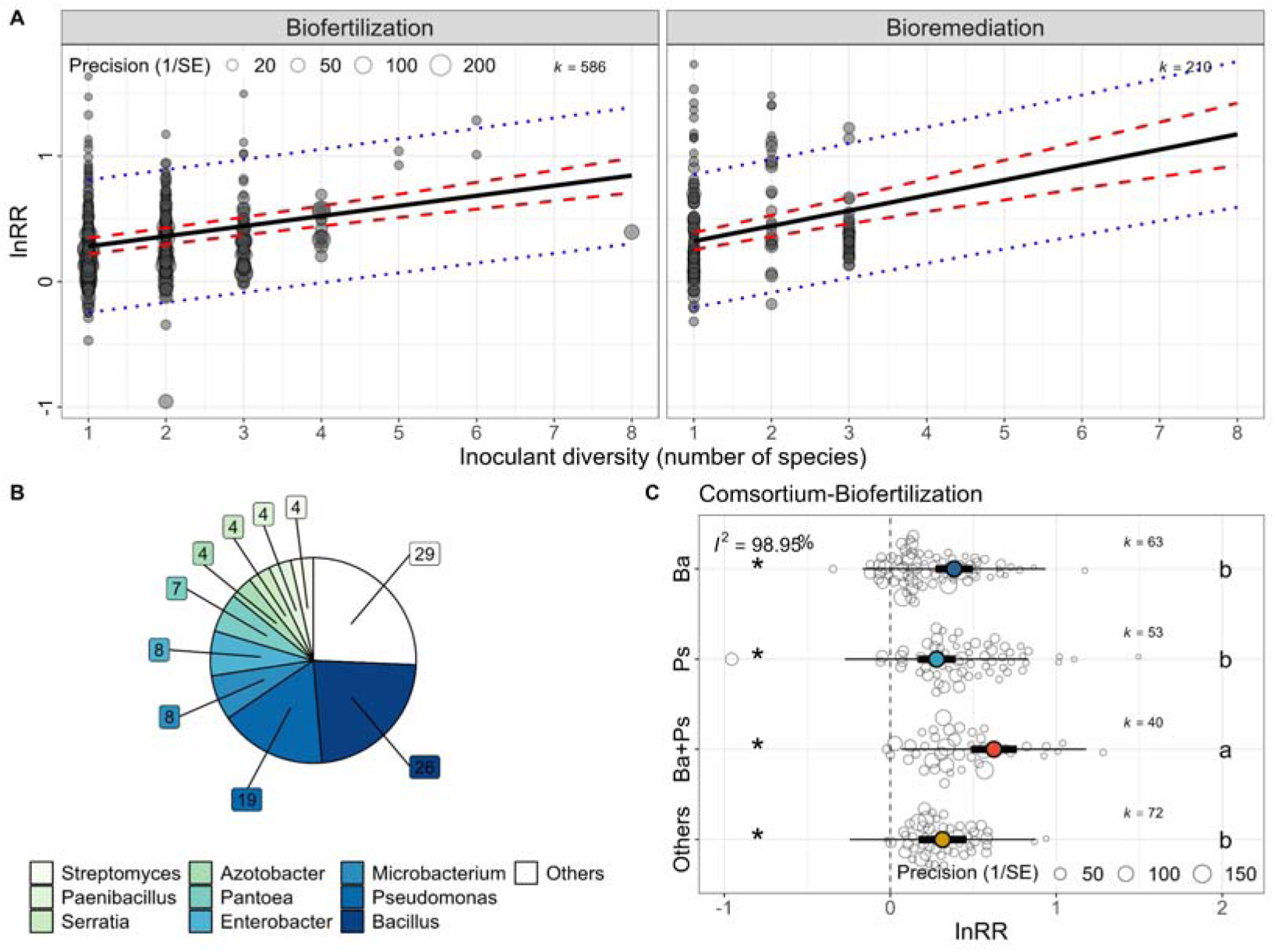
Synergistic effect within consortium contributes to the inoculation effect. (A) Meta-regression analyses show the relationship between effect size (lnRR) and inoculant diversity (estimates = 0.08 and 0.12, p-value < 0.0001 and 0.0001 for biofertilization and bioremediation, respectively). The red and blue dashed lines represent 95% confidence and 95% credibility intervals, respectively. (B) The number of studies using different bacteria (genus) for consortium construction. (C) The comparison of effect sizes between consortia containing *Bacillus* (Ba), *Pseudomonas* (Ps), *Bacillus*+*Pseudomonas* (Ba+Ps), and Others indicating the synergistic effect between *Bacillus* and *Pseudomonas*. Asterisks denote the rejection of the hypothesis that the estimated mean equals 0 (*p* < 0.001), indicating a significant effect compared to non-inoculant treatments. Different letters on the right suggest a significant difference between subgroups (*p* < 0.05, Tukey’s HSD).

To explore the potential synergistic effect of inoculants, we first summarize the type of inoculants (genus level) used in constructing consortia. The two frequently used inoculants were *Bacillus* and *Pseudomonas,* used in 26 and 19 of 51 studies (Fig. 7B). Compared to these genera, other species were relatively less studied, such as *Microbacterium* (8), *Enterobacter* (8), *Pantoea* (7), and *Azotobacter* (4). We further explored the difference between applying *Bacillus*/*Pseudomonas* and other species in the consortium from our biofertilization dataset (the bioremediation dataset was excluded for this analysis due to limited sample size). The result showed no significant difference in effect sizes among consortia containing *Bacillus* (Ba, lnRR = 0.38, 46.9%, 95% CI = 0.27-0.50), *Pseudomonas* (Ps, lnRR = 0.28, 32.2%, 95% CI = 0.17-0.39) and others (Others, lnRR = 0.31, 36.9%, 95% CI = 0.17-0.66) (Fig. 7C and Table S15). However, the combination of *Bacillus* and *Pseudomonas* in the same consortium (Ba+Ps, lnRR = 0.62, 86.6%, 95% CI = 0.48-0.76) had a significantly higher effect (*p* < 0.01) compared to non-combined utilizations (lnRR = 0.38 and 0.28, for *Bacillus* and *Pseudomonas*, respectively) (Fig. 7C and Table S15).

## 4. Discussion

### 4.1. The use of bacterial consortia improves the inoculation effect when compared to single species

In the present meta-analysis, we estimated the overall effect sizes of bacterial inoculation for biofertilization and bioremediation and compared the difference between single-species and consortium inoculants. Only experiments conducted in living soils were included to obtain quantitative results close to natural conditions. Our results showed that bacterial inoculants, in isolation or in a consortium, significantly promoted the functions (i.e., biofertilization and bioremediation) there were selective for when introduced into living soils. For example, applying single species showed an increased effect on plant growth promotion on average by 29%, which is similar to that reported in a recent meta-analysis (J. Li et al., 2022), which suggests the yield increase (32% on average) after the application of microbial inoculants. Importantly, compared to single-species inoculation, we found that consortium applications had a significantly larger estimated effect size for biofertilization (48% on average). Such difference was also observed for bioremediation (increased by 80% and 48% in consortium and single-species treatments, respectively).

The reduced plant growth variation after either consortium or single-species applications in pot/greenhouse conditions implies that using inoculants promotes the uniformization of plant growth over time. For instance, genotypic differences in the plant itself are more likely to dominate its growth traits in the control treatment, while after the application of the inoculant, the plant directly benefits from better conditions in the inoculated soil and thus achieves more stable traits, just as chemical fertilization weakens the phenotypic traits among different plant genotypes (Rengel and Marschner, 2005; Gomez-Coronado et al., 2016). However, in the context of bioremediation, the variation of pollutant removal did not significantly decrease after inoculant applications. This might be due to the lower heterogeneity of the pollution remediation system, in which beneficial bacteria interacted directly with pollutants. Together, these results provide strong empirical evidence that inoculating bacterial consortia in the soil promotes plant growth and pollutant removal more efficiently than using single inoculants and that inoculation (single or in a consortium) improves the stability of plant growth.

### 4.2. Inoculation effect influenced by different regulators

Our second hypothesis that environmental conditions influence the inoculation effect was supported by our results. We found that the experimental condition is an essential factor determining the success of the inoculation, where a significantly larger effect size the decrease in the plant growth variation were observed in the pot/greenhouse experiment after using consortia for biofertilization compared with that in the field condition. For bioremediation, there was no significance between pot/greenhouse and microcosm experiments, probably due to the relatively stable environmental conditions in the soils where the pollutants are present. These results were in line with the previous findings of a meta-analysis (Rubin et al., 2017) and highlight the importance of environmental conditions. For example, it has been reported that PGPR isolates could promote chickpea growth in controlled conditions but failed in a natural setting (Laabas et al., 2017). Despite this, our results showed that a better inoculation effect was achieved in consortium treatments in relation to single-species treatments in both pot/greenhouse and field conditions. Therefore, consortium inoculation can perform better than single strains, although its efficacy may be diminished in the field compared to greenhouse results.

In terms of inoculation methods, we found no significant differences in effect sizes between soil inoculation and seed/seedling inoculation either for single-species or consortium treatments, which was consistent with the results reported by meta-analysis focusing on single-species inoculations (Rubin et al., 2017). However, this does not mean that the inoculation method can be freely selected. On the contrary, it is highly possible that the appropriate protocol had been determined when each study was carried out, as the protocol for soil inoculation depends on the inoculation purpose and the type of inoculants (Fukami et al., 2016; O’Callaghan, 2016). Besides, another recent meta-analysis suggested that combining multiple inoculation approaches (e.g., soil, seed, and foliar) and biological effectors, such as amino acids and microorganisms, can achieve a better effect (Herrmann et al., 2022). Therefore, developing microbial consortia that can successfully colonize various ecological niches, such as bulk soil, rhizosphere, and even endosphere, may increase the scope and effect of inoculants.

Plant type was evaluated as an essential moderator of the inoculum’s function for biofertilization. This might be related to the different objectives of each estimation, which are grain yield for cereal plants and total biomass for non-cereal plants. This may lead to significantly lower effects of cereal crops than non-cereal plants. In contrast, the inoculation effect did not rely on pollutant types, probably due to the appropriate strains had been selected when performing inoculation.

In the context of living soil, inoculated species often suffer competition and suppression from the native soil microbiome, which could result in lower survival over time and a weak inoculation effect (Mallon et al., 2015; O’Callaghan et al., 2022; Liu and Salles, 2023). Therefore, assessing whether the inoculation effect relies on the experiment duration is important while the beneficial effect may decrease due to the decay or extinct of inoculants. We found a higher regression estimate under consortium treatment than single-species treatment for bioremediation, implying the higher efficacy of consortium inoculation. However, our results showed that the regression model’s estimates are close to 0 for both biofertilization and bioremediation, which suggests that the correlation between effect sizes and experiment durations should not be strongly interpreted. Nonetheless, these findings imply that microbial inoculants can have beneficial effects at different time scales and may reflect the successful establishment of inoculants. Indeed, it was reported that the inoculation effect might increase with time as the inoculants are successfully established in the recipient soil (Neuenkamp et al., 2019). However, since studies rarely reported on the survival of inoculants in soil, it is still difficult to reveal how inoculants’ survival correlates with their beneficial effects.

Assessing the impact pattern of original soil properties on the inoculation effect potentially provides instruction for the inoculation strategy. We observed the impact pattern of soil pH on lnRR depending on utilization purposes (i.e., biofertilization and bioremediation). The appropriate soil pH is neutral (pH = 6-8) for biofertilization, resulting in a relatively high inoculation effect, which may be due to the poor survival of inoculants in the relatively extreme pH environment. Soil pH is acknowledged as an important factor for bioremediation and the high-pH environment often lead to lower availability of heavy metals in soil (Neina, 2019). This could be the reason that the observed negative linear regression between pH and lnRR for bioremediation in this study, suggesting the lower inoculation effect of pollutants removal in soil with high pH. Therefore, applying bacterial inoculants in appropriate soil (pH = 6-8) or use other technologies to adjust soil pH may achieve a better inoculation effect. Regarding soil organic matter and available N and P, the observed the significant and positive correlation between their contents and lnRR for biofertilization indicates that providing a soil environment that is relatively rich in nutrients is conducive to the function of inoculated agents. Indeed, a large number of studies have shown that measures such as organic fertilization can promote crop yields by regulating the bioavailability of soil nutrients and functional microorganisms (e.g., P-solubilizing bacteria) (Bi et al., 2020; K. Li et al., 2022). In short, applying microbial inoculants can be supplemented by other management measures such as fertilization to amplify the beneficial inoculation effect in the field.

### 4.3. Synergistic effect within consortium contributes to the inoculation effect

Promoting facilitative interactions and complementary functions between inoculants within a consortium is expected to be desirable for stable performance over a prolonged cultivation (Jiménez et al., 2017; Puentes-Téllez and Falcao Salles, 2018). Our results demonstrate that the inoculant diversity significantly and positively contributes to the inoculation effect. This may explain the advantage of consortia applications over single strains for both biofertilization and bioremediation in living soils, regardless of various conditions. The possible reason is that not only the functions of different inoculants under culture conditions were verified when the communities were constructed in each study, but also the functional complementarity of different microorganisms was considered to optimize the inoculum effect (Tang et al., 2020; Kumawat et al., 2021; Sun et al., 2022).

Indeed, we further demonstrate our third hypothesis that the synergistic effect between inoculants (i.e., *Bacillus* and *Pseudomonas*) within a consortium occurred in living soil by comparing the effects sizes among four groups (applying consortia containing *Bacillus*, *Pseudomonas*, *Bacillus*+*Pseudomonas*, and others). The highest effect size was observed when consortia were constructed by combining *Bacillus* and *Pseudomonas*, which increased plant growth by 86.6%. These two taxa were reported to be frequently used as microbial inoculants for biofertilization and bioremediation (Santoyo et al., 2021; J. Li et al., 2022; Wu et al., 2022) and may function in a complementary fashion in the soil. *Bacillus* is a spore-forming bacteria that can positively affect plant growth through the excretion of auxin or cytokinin-type compounds. At the same time, *Pseudomonas* often exhibit a great potential to inhibit pathogens and diseases due to their rapid growth and antibiotics synthesis, thereby good colonization and competition for niches in the rhizosphere (Santoyo et al., 2012). Moreover, a recent study showed that inoculated *Bacillus* could recruit indigenous *Pseudomonas* through synergistic biofilm formation and syntrophic cooperation to enhance plant-growth-promoting and salt stress-relieving ability (Sun et al., 2022), which suggests a direct interaction between them leading to the synergistic effect. This synergistic effect between *Bacillus* and *Pseudomonas* may also exist in the context of bioremediation approaches (Aravinthan et al., 2016; Fakhar et al., 2022). Thus, our results pointed out an important direction of complementarity or synergism between species when constructing and applying microbial consortia in the soil. However, it is not possible to estimate the synergistic effect between other taxa in this study (owing to limited data) and explore the efficacy of different patterns underlying such synergistic effect with the meta-analyses, as most of the studies do not clarify the mechanisms driving the selection of strains for the consortium. For example, the common method is to select some strains with obvious growth-promoting or pollutant-degrading characteristics for combination (Visioli et al., 2015; Kumar et al., 2017, 2021), but the interactions between these strains are rarely verified and reported. Studies on these are thus needed in the future.

Our meta-analysis demonstrates a significant advantage of applying bacterial consortia on the estimated beneficial effect under various conditions. Despite this, the heterogeneity among effect sizes in all models was high, indicating that there are large inconsistencies in the effect of inoculations (see effect size distribution from Figures and heterogeneity from Supplementary Tables). Such inconsistency implies excellent potential but a significant risk in microbial inoculants’ development and commercial application. Next, we discuss the perspectives for constructing and inoculating microbial consortia.

### 4.4. Perspectives for constructing and inoculating microbial consortia

*Greenhouse ≠ Field, Soil A ≠ Soil B.* Our result shows that the inoculation effect of the bacterial consortium is primarily regulated by environmental conditions, with more pronounced plant-growth promotion and reduced variability of inoculation effect observed in greenhouse experiments. This indicates that the inoculation effect can be severely weakened in the field condition, possibly due to complex environmental disturbances. Therefore, this suggests that the results of greenhouse experiments cannot fully reflect the inoculation effect in field conditions and serve as favorable evidence to support the commercial application of microbial consortia. Thus, we advocate that robust experiments in field conditions are necessary. Moreover, the physiochemical properties of the soil itself need to be considered. For example, regulating soil pH to neutrality, and increasing soil organic matter, N, and P content can improve the inoculation effect. Furthermore, experiments including environmental variation can be used to test the vulnerability of microbial inoculants, however, such studies are rarely done.

*A consortium is better than a single strain.* The current result encourages the application of microbial consortia in living soil for improved biofertilization and bioremediation, regardless of various conditions. Constructing consortia relies on understanding the functionality of different single strains and the potential interactions of different inoculants within a consortium. Selecting microorganisms with complementary and compatible functions that can assist colonization and promote the function of others could be the principle of consortium constructions. Since more species in a consortium may increase its multifunctionality and stability (Jiménez et al., 2018), we propose to (i) construct microbial consortia combining metabolic complementarity and other functional traits such as functional diversity, composition, and redundancy; (ii) systematically obtain inoculation impacts on recipient soils and/or plants. On one hand, microbial species with dynamic metabolic strategies, meaning the ability to diversify their consumption of different resources, are more likely to coexist within a consortium (Pacciani-Mori et al., 2020). On the other hand, if there is a larger niche overlap between the inoculated consortium and the recipient community, it may result in a more severe disturbance in the recipient community and lower survival of inoculants, particularly when resource competition drives the coalescence of two communities (Liu and Salles, 2023). In addition, temporal differences in interactions and synergistic effects should also be considered, for example by manipulating the consortia composition to manipulate different microorganisms to function across time scales (Jiménez et al., 2018). Furthermore, cocktails of inoculants may include beneficial AMF (Verbruggen et al., 2013), bacteriophages (Balogh et al., 2010; Pratama et al., 2020), and protozoa (Kachieng’a and Momba, 2018) towards the multi-directional steering of soil ecosystem functions.

*Is legacy a benefit or a risk?* Microbial inoculants often leave a footprint in the recipient soil (Mawarda et al., 2020; Cornell et al., 2021), while such legacy effect in reshaping soil microbiome is largely overlooked (Liu et al., 2022). Although studies with single-strain and consortium inoculations have shown that the effects of inoculum on soil microbial communities can be significant, at least in the short term (e.g., tens of days) (Mallon et al., 2018; Wang et al., 2018; Xing et al., 2021), the long-term resilience of microbial community composition and the mechanism of its resistance against inoculations remain unknown. The following question is whether there is any potential risk to agroecosystems once the soil native community is altered for a long time. For example, is the surviving inoculant pathogenic to other plants or contributing to plant susceptibility, and will the subsequent crop be able to adapt to the altered indigenous community? For example, the case has been reported that the inoculated AMF induced rice infection to several antagonists in both field and greenhouse conditions (Bernaola et al., 2018). The other side of the coin is that we may engineer soil-native communities through inoculation to develop technologies and strategies for tackling global challenges in the climate change context (Tsoi et al., 2019; Liu et al., 2022). Moreover, the ecological restoration of degraded soils might be achieved by introducing various soil microbial communities (Calderón et al., 2017). Notably, the benefits and potential risks after inoculation should be measured to strike a balance between the two (Liu et al., 2022).

## 4. Conclusions

This meta-analysis provides empirical evidence supporting the advantages of bacterial consortia over single strains for improving plant growth and pollutant removal in living soil. Such advantages were rooted in (i) an increased mean effect size but decreased variation of plant growth; (ii) an increased mean effect size of pollutant removal; and (iii) a stable better effect under various conditions (e.g., environmental conditions, inoculation methods, plant and pollutant types, inoculation durations, and original soil properties). The mechanism of the better inoculation effect of bacterial consortia was characterized as the positive correlation between inoculant diversity and effect sizes for both biofertilization and bioremediation and revealed as the synergistic effect among members (e.g., *Bacillus* and *Pseudomonas*) within consortia. Increasing soil organic matter, N, and P content and regulating soil pH to 6-7 are beneficial to inoculation effects. Overall, our results show great promise of microbial consortia for biofertilization and bioremediation in living soils. Although some of these results were expected and unraveling the soil biological processes that influence inoculation efficacy remains difficult, this meta-analysis provides the first proper quantification of the effect sizes, supported by statistical analysis across multiple experiments. It is foreseeable that ideal microbial consortia are likely diverse, and their empirical inoculation effects are context-dependent. The systematic framework for developing consortia should be established in the future, and robust experiments for validations should be performed before large-scale applications to achieve environmentally friendly ecosystem management.

## Author contributions

X.L. and J.F.S collaborated on conceptualization. X.L. conducted data collection and analysis and manuscript writing. S.M. helped in data collection and analyses. All authors helped review and revise the manuscript and agreed to the current version of the manuscript.

## Data availability

All data and code for running the analysis and data visualization are available from OSF: https://osf.io/n3zt9/?view_only=f5a1dfdbc20a4d159af49117eb91b681.

## Acknowledgments

This work was financed by the ERA-NET Cofund SusCrop project potatoMETAbiome (Grant No 771134). X.L. and S.M. were supported by scholarships from the China Scholarship Council.

## Appendix A. Supplementary data

The Supplementary information includes Supplementary Fig. S1-5 and Tables S1-15.

The list of all references involved in the meta-analysis is available from OSF: https://osf.io/n3zt9/?view_only=f5a1dfdbc20a4d159af49117eb91b681.

