## Supplemental Materials for "Do inoculated microbial consortia perform better than single strains in living soil? A meta-analysis"

The Supplementary information includes Supplementary Figures S1-5 and Tables S1-15.

**Supplementary Figures**

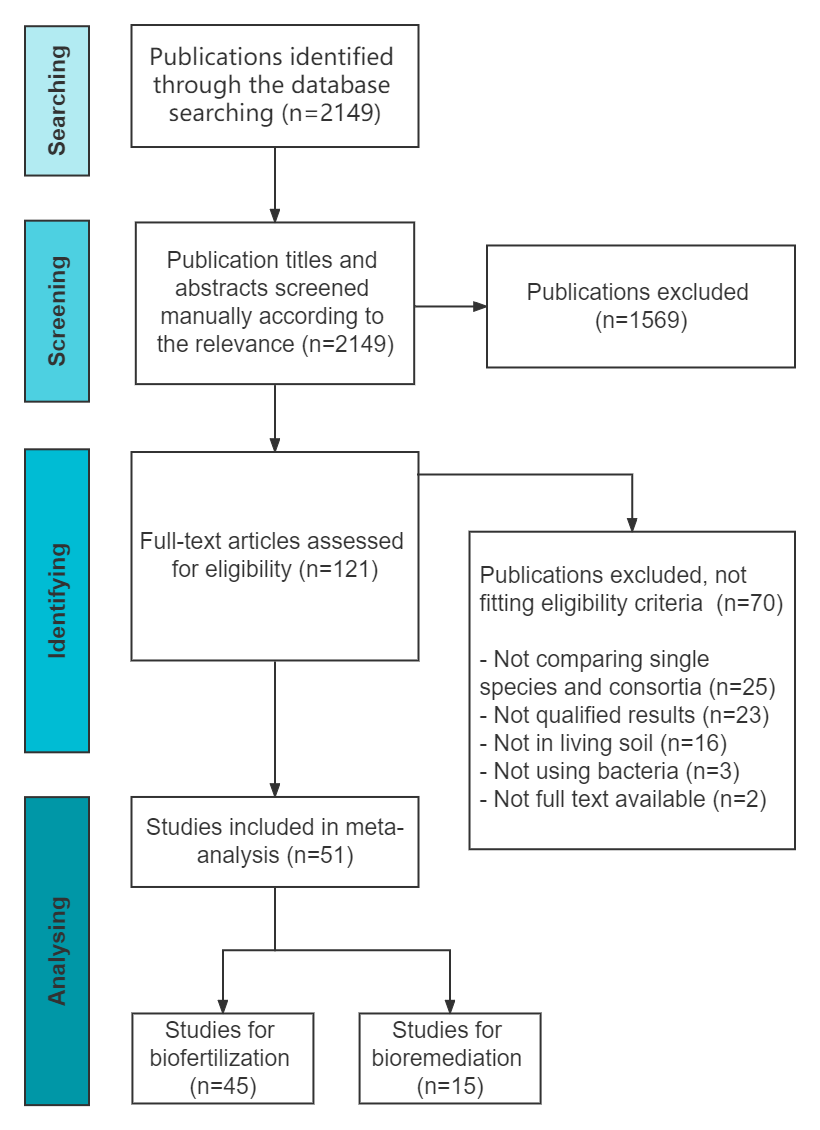

Fig. S1. The PRISMA diagram shows the workflow and the number of studies included in each step of the meta-analysis. The studies were excluded following the eligibility criteria, such as studies using only single-species or consortia, performed in sterile or non-soil substrates, and in which the results could not be extracted directly from the Figures (see Methods for details).

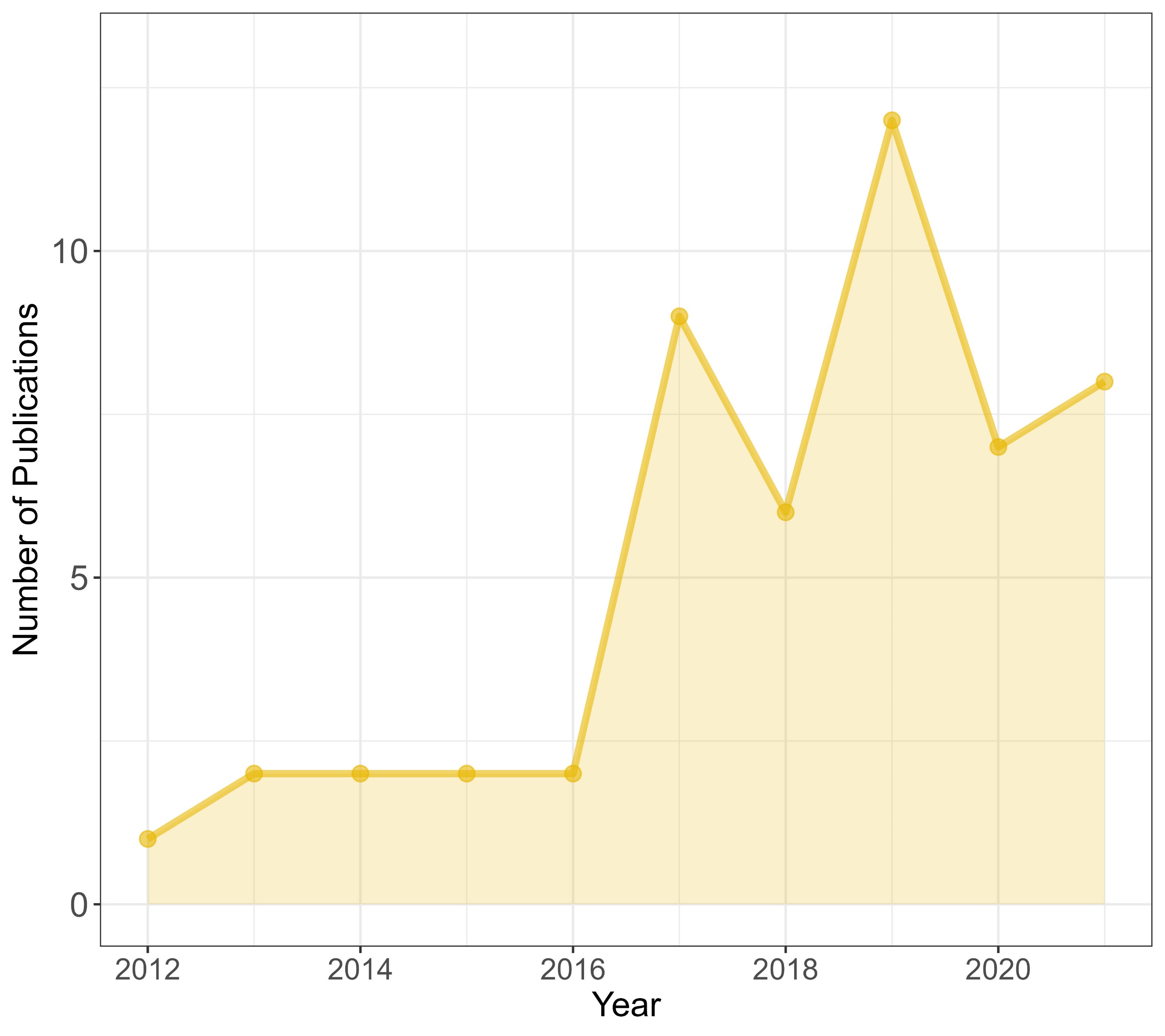

Fig. S2. The publication time of 51 studies.

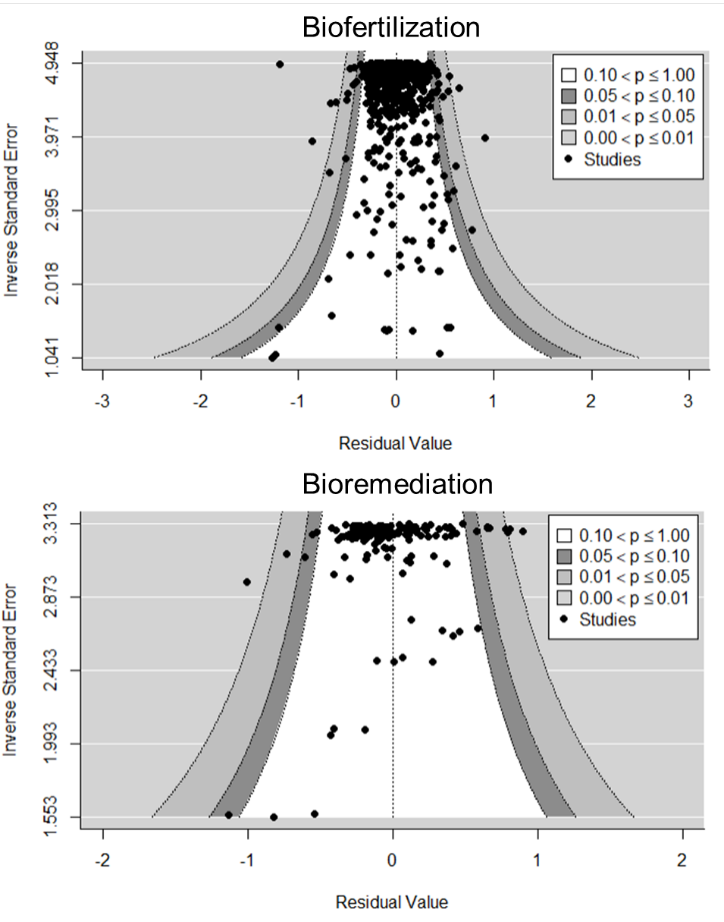

Fig. S3. Funnel plots testing for asymmetry for effect sizes from biofertilization and bioremediation datasets. The conventional Egger’s regression test for publication bias was conducted and shown in Table S1. No significant publication bias was found in our datasets.

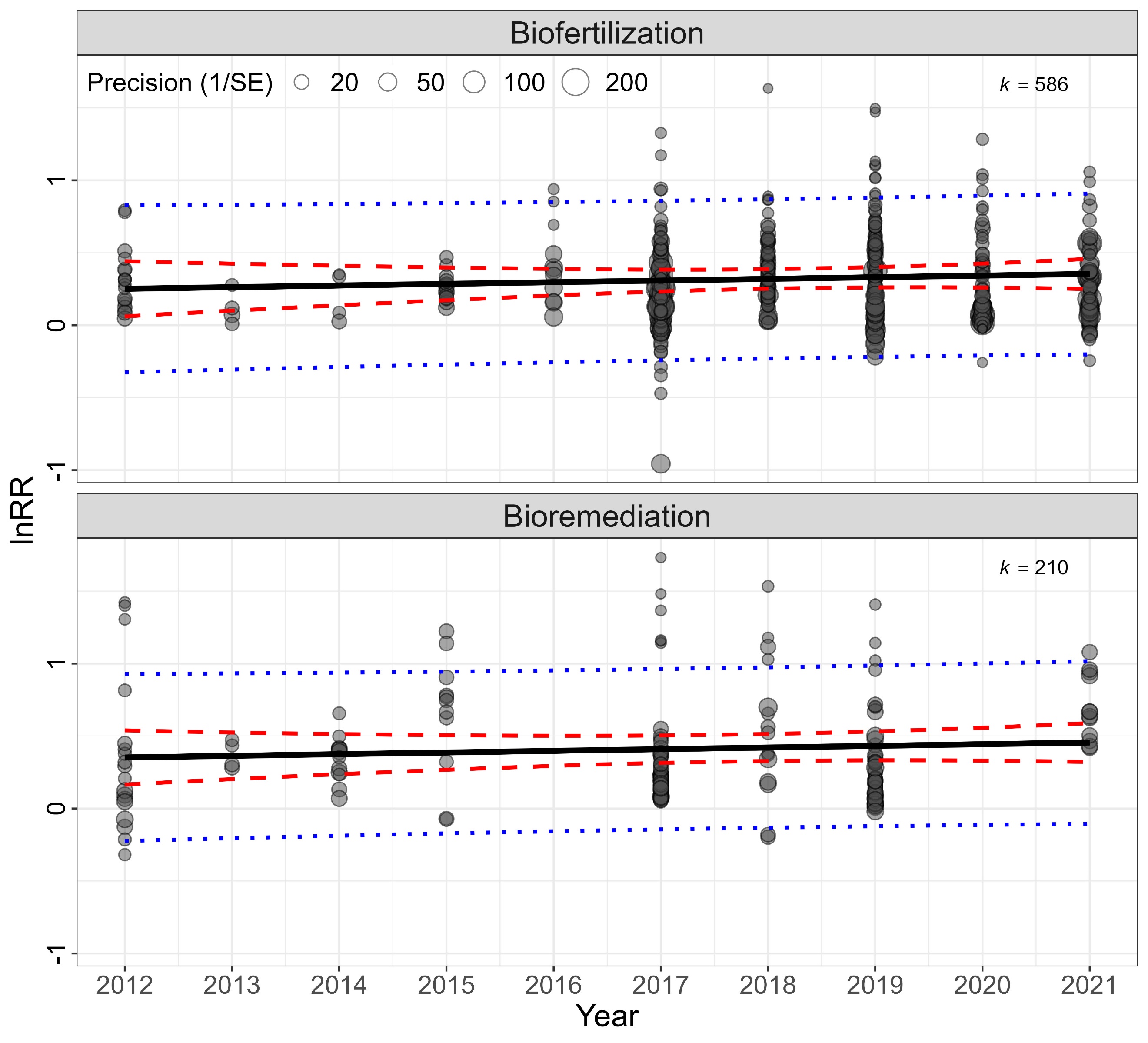

Fig. S4. Meta-regression model for assessing the time-lag bias. No significant time-lag tendency was observed both for biofertilization and bioremediation datasets. The red and blue dashed lines represent 95% confidence and 95% credibility intervals, respectively. The statistical results are shown in Table S2.

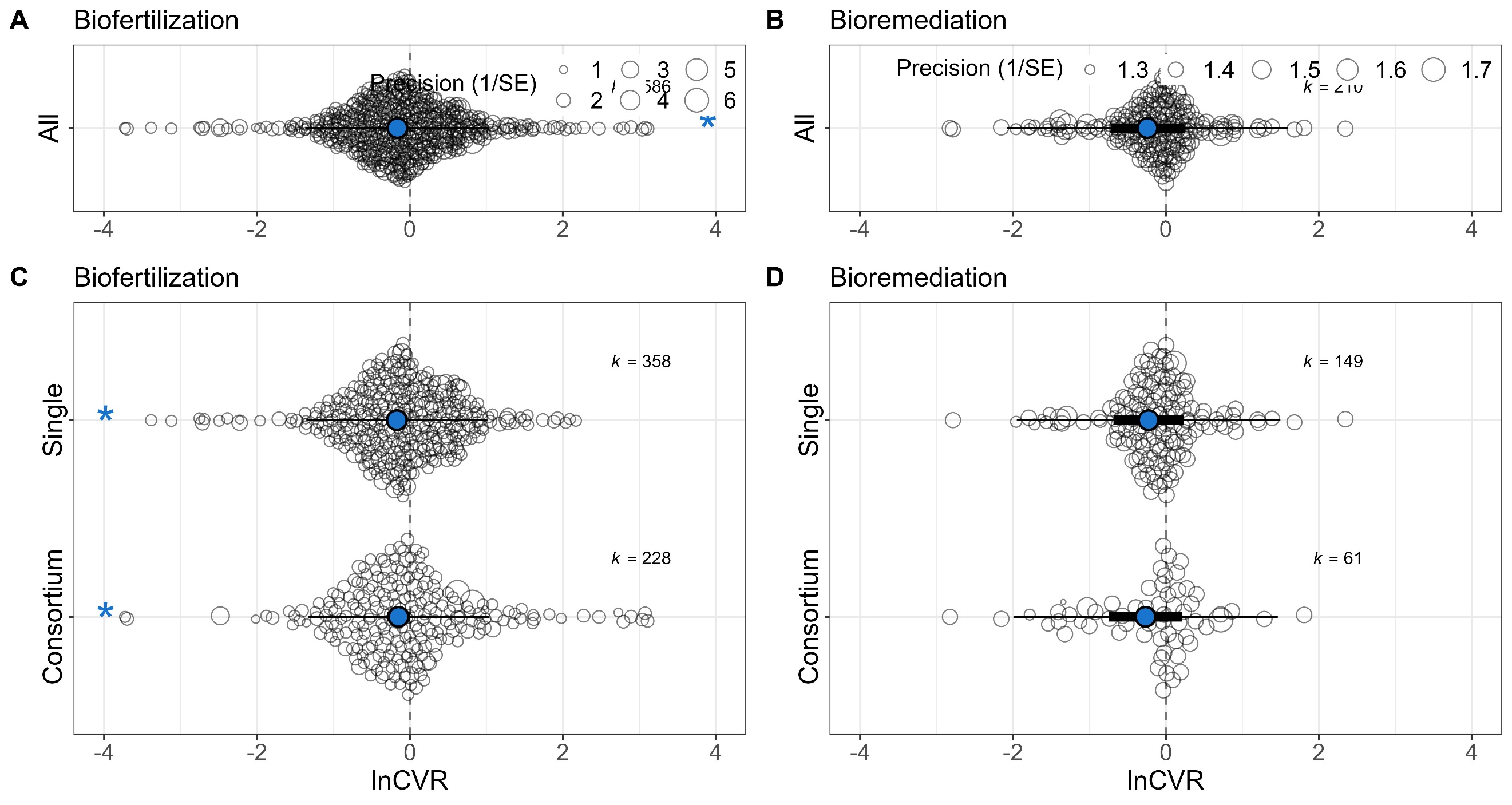

Fig. S5. The variability (lnCVR) of beneficial effect for biofertilization and bioremediation functions. The overall effects (A, B) and comparisons between single-species and consortium inoculants (C, D) were estimated. Central points represent estimated means, thick bars represent 95% confidence intervals, and thin bars represent 95% prediction intervals. Asterisks denote the rejection of the hypothesis that the estimated mean equals 0 (*p* < 0.001), indicating a significant effect compared to non-inoculant treatments.

**Supplementary Tables**

Table S1. The conventional Egger’s regression test for publication bias. A *p*-value greater than 0.05 rejects the hypothesis of data asymmetry in the funnel plot.

| Data | Test | Estimate | t | df | *p* |
| --- | --- | --- | --- | --- | --- |
| Biofertilization | Asymmetry | 0.18 (0.16-0.20) | 1.48 | 584 | 0.14 |
| Bioremediation | Asymmetry | 0.22 (0.13-0.32) | 1.16 | 208 | 0.25 |

Table S2. Meta-regression model for assessing the time-lag bias. A *p*-value greater than 0.05 suggests no significant correlation between publication year and effect size, indicating no obvious time-lag bias. CL represents confidence interval.

| Model | Estimate | LowerCL | UpperCL | t-value | p-value |
| --- | --- | --- | --- | --- | --- |
| Biofertilization (Publication year) | 0.0114 | -0.0163 | 0.0391 | -0.1836 | 0.4194 |
| Bioremediation (Publication year) | 0. 0114 | -0.0166 | 0. 0395 | -0.3199 | 0.4194 |

Table S3. Overall effect of bacterial inoculants for log response ratio (lnRR) compared to the non-inoculant treatment. CL and PR represent confidence interval and prediction interval, respectively.

| Model | Estimate | LowerCL | UpperCL | LowerPR | UpperPR |
| --- | --- | --- | --- | --- | --- |
| Biofertilization  (All_lnRR) | 0.3059  (35.78%) | 0.2447  (27.72%) | 0.3671  (44.35%) | -0.1836  (-16.77%) | 0.7954  (121.53%) |
| Bioremediation  (All_lnRR) | 0.4572  (57.96%) | 0.2834  (32.76%) | 0.6310  (87.95%) | -0.3199  (-37.70%) | 1.2343  (243.60%) |

Table S4. The effects of subgroups (single inoculant and consortium) for log response ratio (lnRR) compared to the non-inoculant treatment. CL and PR represent confidence interval and prediction interval, respectively.

| Model | Estimate | LowerCL | UpperCL | LowerPR | UpperPR |
| --- | --- | --- | --- | --- | --- |
| Biofertilization (Consortium_lnRR) | 0.3957 (48.54%) | 0.3333 (39.56%) | 0.4580 (58.09%) | -0.0618 (-5.99%) | 0.8531 (134.69%) |
| Biofertilization (Single_ lnRR) | 0.2518 (28.64%) | 0.1916 (21.12%) | 0.3119 (36.60%) | -0.2054 (-18.57%) | 0.7089 (103.18%) |
| Bioremediation  (Consortium_lnRR) | 0.5904 (80.47%) | 0.4187 (52.00%) | 0.7621 (114.28%) | -0.1173 (-11.07%) | 1.2981 (266.23%) |
| Bioremediation  (Single _ lnRR) | 0.3893 (47.59%) | 0.2234 (25.03%) | 0.5552 (74.23%) | -0.3170 (-27.17%) | 1.0956 (199.10%) |

I^2^ = 99.15%, R^2^_marginal = 0.07, R^2^_conditional = 0.76.

Table S5. Overall effect of bacterial inoculants for variation in inoculation impacts (lnCVR) compared to the non-inoculant treatment. CL and PR represent confidence interval and prediction interval, respectively.

| Model | Estimate | LowerCL | UpperCL | LowerPR | UpperPR |
| --- | --- | --- | --- | --- | --- |
| Biofertilization  (All_ lnCVR) | -0.1621  (-14.96%) | -0.2831  (-24.66%) | -0.0411  (-4.03%) | -1.3674  (-292.51%) | 1.0432  (183.83%) |
| Bioremediation  (All_ lnCVR) | -0.2400  (27.12%) | -0.7233  (-106.12%) | 0.2434  (27.56%) | -2.0772  (-87.47%) | 1.5973  (393.97%) |

Table S6. The effects of subgroups (single inoculant and consortium) for variation in inoculation impacts (lnCVR) compared to the non-inoculant treatment. CL and PR represent confidence interval and prediction interval, respectively.

| Model | Estimate | LowerCL | UpperCL | LowerPR | UpperPR |
| --- | --- | --- | --- | --- | --- |
| Biofertilization (Consortium_ lnCVR) | -0.1486 (-16.02%) | -0.2956 (-34.39%) | -0.0016 (-0.16%) | -1.3345 (-279.81%) | 1.0373 (182.16%) |
| Biofertilization (Single_ lnCVR) | -0.1701 (-18.54%) | -0.2998 (-34.96%) | -0.0404 (-4.12%) | -1.3540 (-287.29%) | 1.0138 (175.61%) |
| Bioremediation  (Consortium_ lnCVR) | -0.2664 (-30.53%) | -0.7428 (-110.18%) | 0.2100 (23.37%) | -1.9957 (-635.74%) | 1.4629 (331.85%) |
| Bioremediation  (Single _ lnCVR) | -0.2268 (-25.46%) | -0.6860 (-98.58%) | 0.2324 (26.16%) | -1.9515 (-603.92%) | 1.4979 (347.23%) |

Table S7. The impact of experiment types on effect sizes for biofertilization. The experiment type indicates whether the experiment was conducted in soil microcosm (without plants), pot/greenhouse (with plant), or filed (with plant) conditions. CL and PR represent confidence interval and prediction interval, respectively.

| Name | Condition | Estimate | LowerCL | UpperCL | LowerPR | UpperPR |
| --- | --- | --- | --- | --- | --- | --- |
| Pot | Single | 0.2652  (30.37%) | 0.2036  (22.58%) | 0.3268  (38.65%) | -0.1866  (-17.02%) | 0.7170  (104.83%) |
| Field | Single | 0.1912  (21.07%) | 0.0993  (10.44%) | 0.2831  (32.72%) | -0.2657  (-23.33%) | 0.6482  (91.21%) |
| Pot | Consortium | 0.4164  (51.65%) | 0.3518  (42.16%) | 0.4811  (61.79%) | -0.0358  (-3.52%) | 0.8687  (138.38%) |
| Field | Consortium | 0.3198  (37.69%) | 0.2280  (25.61%) | 0.4116  (50.92%) | -0.1371  (14.69%) | 0.7767  (117.43%) |

I^2^ = 98.77%, R^2^_ marginal = 0.10, R^2^_ conditional = 0.75.

Table S8. The impact of experiment types on effect sizes for bioremediation. The experiment type indicated whether the experiment was conducted in soil microcosm (without plants), pot/greenhouse (with plant), and filed (with plant) conditions. CL and PR represent confidence interval and prediction interval, respectively.

| Name | Condition | Estimate | LowerCL | UpperCL | LowerPR | UpperPR |
| --- | --- | --- | --- | --- | --- | --- |
| Pot | Single | 0.3951  (48.45%) | 0.2032  (22.53%) | 0.5870  (79.86%) | -0.3417  (-28.94%) | 1.1319  (210.15%) |
| Microcosm | Single | 0.3880  (47.40%) | -0.0111  (-1.10%) | 0.7871  (119.70%) | -0.4277  (-34.80%) | 1.2037  (233.24%) |
| Pot | Consortium | 0.6059  (83.29%) | 0.4077  (50.34%) | 0.8040  (123.45%) | -0.1326  (-12.42%) | 1.3443  (283.55%) |
| Microcosm | Consortium | 0.5141  (67.21%) | 0.1008  (10.61%) | 0.9274  (152.79%) | -0.3086  (-26.55%) | 1.3368  (280.68%) |

I^2^ = 98.22%, R^2^_ marginal = 0.07, R^2^_ conditional = 0.88.

Table S9. The impact of inoculation methods on effect sizes for biofertilization. Inoculation methods includes soil inoculation and seed inoculation, which depict the application of microbial inoculants either directly by pouring into the soil or by soaking the seeds/seedlings. CL and PR represent confidence interval and prediction interval, respectively.

| Name | Condition | Estimate | LowerCL | UpperCL | LowerPR | UpperPR |
| --- | --- | --- | --- | --- | --- | --- |
| Soil inoculation | Single | 0.2330  (26.24%) | 0.1597  (17.32%) | 0.3064  (35.85%) | -0.2242  (-20.08%) | 0.6903  (99.43%) |
| Seed inoculation | Single | 0.2918  (33.88%) | 0.1883  (20.72%) | 0.3953  (48.48%) | -0.1712  (-15.73%) | 0.7548  (112.72%) |
| Soil inoculation | Consortium | 0.3618  (43.59%) | 0.2843  (32.88%) | 0.4393  (55.16%) | -0.0961  (-9.16%) | 0.8197  (126.98%) |
| Seed inoculation | Consortium | 0.4532  (57.33%) | 0.3484  (41.68%) | 0.5580  (74.72%) | -0.01012  (-1.01%) | 0.9166  (150.08%) |

I^2^ = 98.79%, R^2^_ marginal = 0.12, R^2^_ conditional = 0.76.

Table S10. The impact of inoculation methods on effect sizes for bioremediation. Inoculation methods includes soil inoculation and seed inoculation, which depict the application of microbial inoculants either directly by pouring into the soil or by soaking the seeds/seedlings. CL and PR represent confidence interval and prediction interval, respectively.

| Name | Condition | Estimate | LowerCL | UpperCL | LowerPR | UpperPR |
| --- | --- | --- | --- | --- | --- | --- |
| Soil inoculation | Single | 0.4113  (50.88%) | 0.2177  (24.32%) | 0.6049  (83.11%) | -0.3265  (-27.86%) | 1.1491  (215.54%) |
| Seed inoculation | Single | 0.3084  (36.12%) | -0.0751  (-7.23%) | 0.6919  (99.75%) | -0.5003  (-39.37%) | 1.1171  (205.60%) |
| Soil inoculation | Consortium | 0.6044  (83.02%) | 0.4042  (49.81%) | 0.8045  (123.56%) | -0.1352  (-12.65%) | 1.3439  (283.40%) |
| Seed inoculation | Consortium | 0.5428  (72.08%) | 0.1467  (15.80%) | 0.9388  (155.69%) | -0.2719  (-23.81%) | 1.3575  (288.65%) |

I^2^ = 98.22%, R^2^_ marginal = 0.07, R^2^_ conditional = 0.88

Table S11. The impact of plant types on effect sizes for biofertilization. The plant type was grouped into cereal and non-cereal plants and the pollutant type was divided into heavy metal and organic pollutant. CL and PR represent confidence interval and prediction interval, respectively.

| Name | Condition | Estimate | LowerCL | UpperCL | LowerPR | UpperPR |
| --- | --- | --- | --- | --- | --- | --- |
| Cereal | Single | 0.1890  (20.80%) | 0.1168  (12.39%) | 0.2612  (29.85%) | -0.2466  (-21.85%) | 0.6246  (86.75%) |
| Non-cereal | Single | 0.3338  (39.63%) | 0.2444  (27.69%) | 0.4232  (52.68%) | -0.1050  (-9.97%) | 0.7727  (116.56%) |
| Cereal | Consortium | 0.3234  (38.18%) | 0.2490  (28.27%) | 0.3978  (48.85%) | -0.1126  (-10.65%) | 0.7594  (113.70%) |
| Non-cereal | Consortium | 0.5088  (66.33%) | 0.4132  (51.16%) | 0.6045  (83.03%) | 0.0687  (7.11%) | 0.9490  (158.31%) |

I^2^ = 98.67%, R^2^_ marginal = 0.16, R^2^_ conditional = 0.75.

Table S12. The impact of pollutant types on effect sizes for bioremediation. The plant type was grouped into cereal and non-cereal plants and the pollutant type was divided into heavy metal and organic pollutant. CL and PR represent confidence interval and prediction interval, respectively.

| Name | Condition | Estimate | LowerCL | UpperCL | LowerPR | UpperPR |
| --- | --- | --- | --- | --- | --- | --- |
| Metal | Single | 0.4238  (52.78%) | 0.2452  (27.79%) | 0.6024  (82.65%) | -0.3126  (-26.85%) | 1.1602  (219.06%) |
| Organic pollutant | Single | 0.3244  (38.32%) | 0.1290  (13.77%) | 0.5198  (68.17%) | -0.4163  (-34.05%) | 1.0651  (190.11%) |
| Metal | Consortium | 0.6226  (86.38%) | 0.4377  (54.91%) | 0.8075  (124.23%) | -0.1154  (-10.90%) | 1.3605  (289.81%) |
| Organic pollutant | Consortium | 0.5351  (70.76%) | 0.3272  (38.71%) | 0.7429  (110.20%) | -0.2090  (-18.86%) | 1.2792  (259.38%) |

I^2^ = 98.31%, R^2^_ marginal = 0.06, R^2^_ conditional = 0.89

Table S13. Meta-regression model for assessing the impact of experimental duration on effect sizes (lnRR). A *p*-value less than 0.05 suggests the significant correlation between experimental duration and effect size. CL represents confidence interval.

| Model | Estimate | LowerCL | UpperCL | t-value | p-value |
| --- | --- | --- | --- | --- | --- |
| Biofertilization (Single-species) | 0.0003 | -0.0003 | 0.0009 | 0.9707 | 0.3322 |
| Biofertilization  (Consortium) | 0.0013 | 0.0001 | 0.0024 | 1.2393 | 0.2163 |
| Bioremediation  (Single-species) | 0.0005 | -0.0003 | 0.0014 | 2.1143 | 0.0271 |
| Bioremediation  (Consortium) | 0.0017 | 0.0003 | 0.0032 | 2.3131 | 0.0214 |

Table S14. Meta-regression model for assessing the impact of inoculant diversity on effect sizes (lnRR). A *p*-value less than 0.05 suggests the significant correlation between inoculant diversity and effect size. CL represents confidence interval.

| Model | Estimate | LowerCL | UpperCL | t-value | p-value |
| --- | --- | --- | --- | --- | --- |
| Biofertilization (Inoculant diversity) | 0.0805 | 0.0624 | 0.0986 | 8.7295 | < 0.0001 |
| Bioremediation (Inoculant diversity) | 0. 1216 | 0.0905 | 0.1526 | 7.6819 | < 0.0001 |

Table S15. The effect sizes of different consortia containing Bacillus (Ba), Pseudomonas (Ps), Bacillus+Pseudomonas (Ba+Ps), and others. CL and PR represent confidence interval and prediction interval, respectively.

| Model | Estimate | LowerCL | UpperCL | LowerPR | UpperPR |
| --- | --- | --- | --- | --- | --- |
| Ba | 0.3845  (46.89%) | 0.2718  (31.23%) | 0.4973  (64.43%) | -0.1669  (-15.37%) | 0.9359  (154.95%) |
| Ps | 0.2790  (32.18%) | 0.1652  (17.96%) | 0.3927  (48.10%) | -0.2726  (-23.86%) | 0.8306  (129.47%) |
| Ba+Ps | 0.6239  (86.62%) | 0.1690  (18.41%) | 0.7620  (114.26%) | 0.0667  (6.90%) | 1.1810  (225.76%) |
| Others | 0.3139  (36.88%) | 0.1690  (18.41%) | 0.4589  (58.23%) | -0.2449  (-21.72%) | 0.8728  (139.36%) |

I^2^ = 99.18%, R^2^_ marginal = 0.16, R^2^_ conditional = 0.81.
